# A neuropsychological basis for temptation-resistant voluntary exercise

**DOI:** 10.1101/2023.07.19.549365

**Authors:** Alexander L. Tesmer, Xinyang Li, Cyra Schmandt, Rafael Polania, Daria Peleg-Raibstein, Denis Burdakov

## Abstract

Despite well-known health benefits of physical activity, many people under-exercise, and what drives prioritization of exercise over other alternatives is unclear. We implement a novel paradigm allowing to study how freely behaving mice rapidly display such prioritizing between time spent on wheel-running and other temptations such as palatable food. Causal manipulations and correlative analyses of underlying appetitive and consummatory psychobehavioral processes revealed this prioritizing to be instantiated by hypothalamic hypocretin/orexin neurons.

## INTRODUCTION

There is an overwhelming agreement in scientific literature and global health guidelines that physical exercise is beneficial for health ^1^. Despite this general knowledge, many people under-exercise, instead overeating highly-palatable food (HPF) that is widely available in many societies ^2-4^. Maintaining exercise despite the presence of HPF temptation would thus be beneficial, however, little is known about how the brain may achieve this. In particular, the neural determinants of arbitration between exercising and HPF consumption are unknown. Hypothalamic orexin neurons (HONs) are a recently characterized neural module that is implicated in regulating food consumption and locomotion over chronic timescales ^5-9^. However, their acute role in eating is unclear ^10, 11^, and whether they are involved in acutely arbitrating between HPF and exercising when multiple choices are available is not known. Here, using voluntary wheel running in mice as a well-established model for human health-beneficial voluntary exercise ^12, 13^, we designed experiments aimed at assessing HONs’ role in temptation-resistant exercise in multiple-choice scenarios that mimic acute decisions in humans of whether to exercise or engage in other pursuits. We then relate these observations to psychological frameworks of appetitive (“approach”) and consummatory (“engagement”) phases of both eating and exercising, and provide a mechanistic neuroeconomic account of preference towards exercise.

## RESULTS

### Temptation-resistant exercise in mice and its link to orexin neurons

Subjects were placed in the center of an arena containing a running wheel (RW) and multiple equidistant alternatives which could freely choose between, thus allowing us to visualize their priorities (Fig. 1A,B). The alternatives, placed in different arms of the maze, included the RW, a novel object, a novel mouse, water, light and dark areas, chow; one arm was either left empty or contained highly-palatable food (HPF, see Methods). When the food option was limited to standard chow, mice chose to split most of their time between RW and chow (grey trace, Fig. 1C). When HPF was added to the alternatives, mice became strongly tempted towards it, substantially reducing their time at chow (colored trace, Fig. 1C). Strikingly, however, RW occupation and usage remained unaltered in the presence of HPF (Fig 1D,E), as did total distance traveled in the arena (Fig. 1F). These observations thus define a mouse model for voluntary exercise-like activity resistant to HPF temptation (henceforth, “TRVE”: **T**emptation-**R**esistant **V**oluntary **E**xercise).

**Fig. 1.**
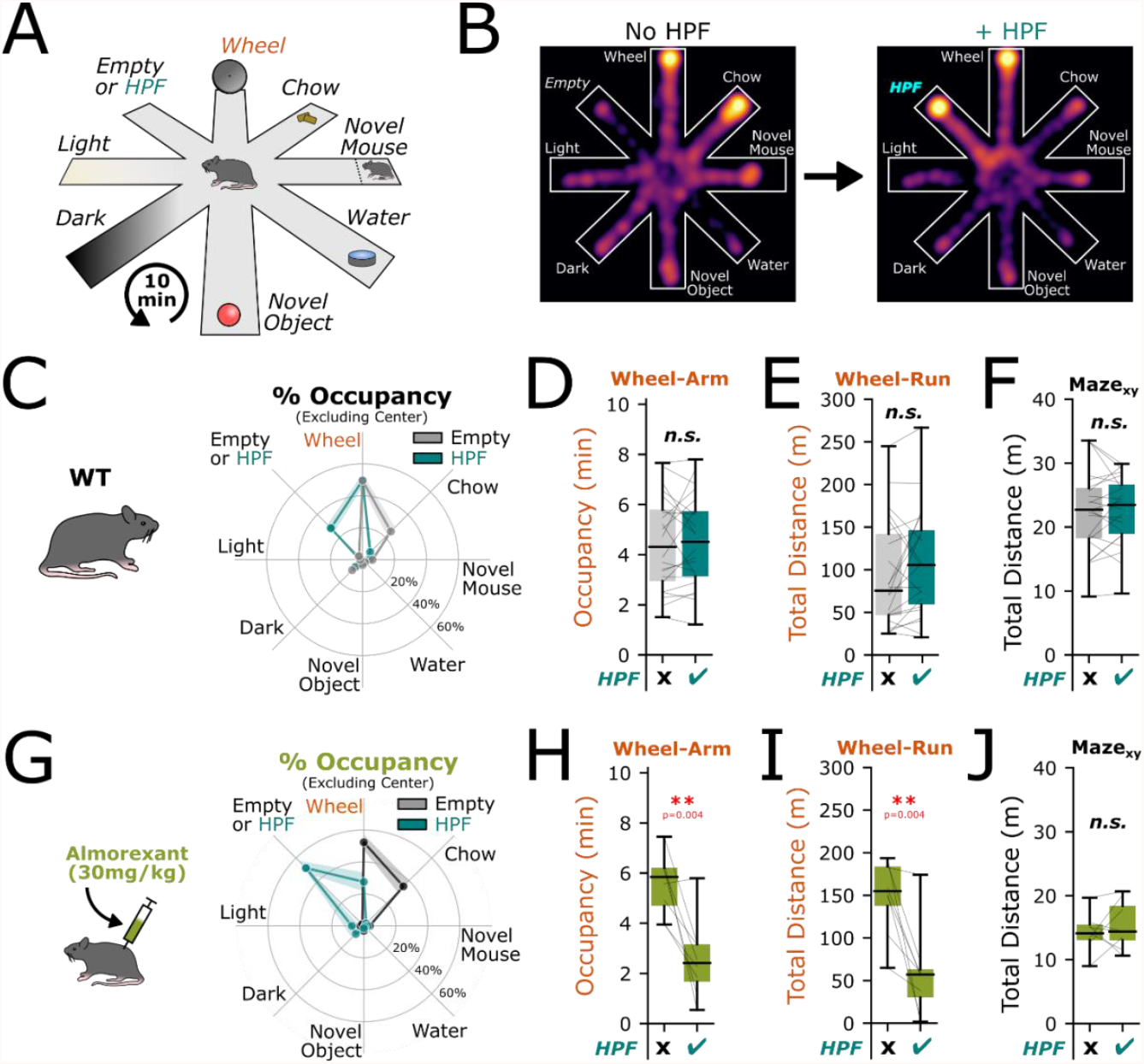
Temptation-resistant voluntary exercise in mice depends on the orexin system. **A** Subjects explored a radial 8-arm arena containing unique alternatives at the end of each arm. Place-preferences of each subject were recorded over a 10-minute period. **B** Heatmaps of an example subject displaying a shift in place preference from the chow-arm (left) towards the HPF, when available (right). **C** Subjects displayed the highest % occupancy in the RW or chow-arm without HPF (grey). Subjects displayed the highest % occupancy in the RW or HPF-arm when the HPF was available (teal). Lines represent mean and SEM of n=22 mice. **D** Effect of HPF on total time spent in the wheel-arm. Two-sided two sample paired t-test, p = 0.804. **E** Effect of HPF on total distance traveled on the RW. Two-sided two sample paired t-test, p = 0.124. **F** Effect of HPF on total distance run in the arena. Two-sided two sample paired t-test, p = 0.858. **G** Mice were injected with 30 mg/kg B.W. almorexant 40 minutes before being placed into the arena. Subjects displayed the highest % occupancy in the RW or chow-arm without HPF (grey). Subjects displayed the highest % occupancy in the RW or HPF-arm when the HPF was available (teal), note that RW-arm % occupancy was reduced when HPF was added. Lines represent mean and SEM of n=8 mice. **H** Effect of HPF on total time spent in the wheel-arm in almorexant-injected subjects. Two-sided two sample paired t-test, p = 0.004. **I** Effect of HPF on total distance traveled on the RW in almorexant-injected subjects. Two-sided two sample paired t-test, p = 0.004. **J** Effect of HPF on total distance run in the arena in almorexant-injected subjects. Two-sided two-sample paired t-test, p = 0.498.

To test whether HONs contribute to TRVE, we suppressed their impact using the orexin receptor antagonist almorexant (ALMO). ALMO abolished TRVE (Fig. 1G) with HPF addition to the maze now significantly reducing the time spent in RW arm (Fig. 1G,H), as well as RW usage (Fig. 2I), while the total distance travelled remained unchanged (Fig. 1J). This indicates that when orexin receptors were antagonized, HPF selectivity reduced RW attraction and usage, rather than general locomotion. Chronic deletion of hypocretin/orexin neurons using the orexin-DTR cell ablation model ^14^, also disrupted TRVE (Supp. Fig S1). Overall, these data indicate that HONs, and orexin receptor signaling in particular, are necessary neural elements for TRVE.

**Fig. 2.**
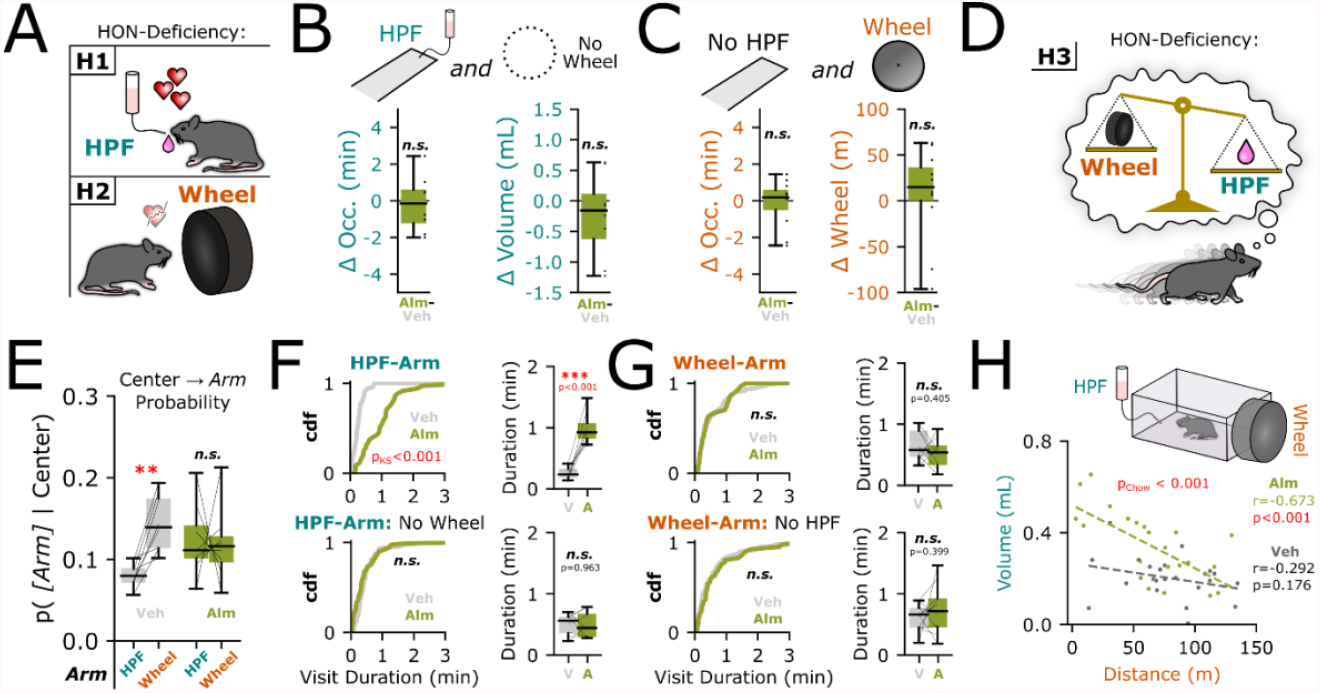
Disentangling psychological processes underlying orexin-dependent TRVE. **A** Two hypotheses explaining the effects of HON-deficiency on TRVE. HON-deficiency may drive mice to increase affinity to the HPF (upper) or decrease affinity to the RW (lower). **B** Almorexant-induced changes in HPF-arm occupancy in an arena without RW available (left, two-sided one sample t-test, p = 0.788, n=8 mice) and HPF consumption (right, two-sided one sample t-test, p = 0.353, n=8 mice). **C** Almorexant-induced changes in RW-arm occupancy in an arena without HPF available (left, two-sided one sample t-test, p = 0.767, n=11 mice) and RW usage (right, two-sided one sample t-test, p = 0.668, n=11 mice). **D** Alternative hypothesis suggesting that HON-deficiency may be explained by a shifted value-comparison between RW and HPF. **E** Probability to enter the HPF or RW arm given that the mouse in the neutral center area. Vehicle-injected mice (grey); two-sided two sample t-test, adjusted p=0.003. Almorexant-injected (green); two-sided two sample t-test, adjusted p=1.000, n = 8 mice. **F** Cumulative distribution plot of all HPF-arm entries from n=8 mice when RW was available after almorexant or vehicle-injected mice (upper left, KS-test p<0.001). Same plot when RW was not available (lower left, KS-test p=0.077). Mouse-averaged visit durations when both RW and HPF were available (upper right, two-sided two sample t-test, p<0.001). Same plot when RW was not available (lower right, two-sided two sample t-test, p=0.963). **G** Cumulative distribution plot of all RW-arm entries from n=8 mice when HPF was available after almorexant or vehicle-injected mice (upper, KS-test p=0.953). Same plot of n = 8 mice when HPF was not available (lower, KS-test p=0.728). Mouse-averaged visit durations when both RW and HPF were available (upper right, two-sided two sample t-test, p=0.405). Same plot when HPF was not available (lower right, two-sided two sample t-test, p=0.399). **H** Mice were placed for 10 minutes into a cage with both HPF and RW available in close proximity. Almorexant and vehicle-injected mice were described by significantly different regression lines. Chow-test, p<0.001. Almorexant-injected mice had a significant linear relationship between HPF and RW usage. Pearson’s r =-0.673, Wald-test, p<0.001. Vehicle-injected mice did not have a significant relationship between HPF and RW usage. Pearson’s r=-0.292, Wald-test p=0.176. In this figure, “appetitive” behaviors are: occ. = spatial occupancy, p=probability of entry, visit duration; “consummatory” behaviors: volume consumed, Wheel = distance covered on RW.

### Orexin impact on behavioral subprocesses related to feeding and exercising

What neurophysiological mechanisms underlie HON-dependent TRVE? It is currently thought that orexin peptides increase locomotion and modulate eating ^5, 15^ (Fig. 2A). However, when HPF but not RW was present, we found that ALMO affected neither appetitive nor consummatory processes related to HPF (here quantified as HPF area occupancy and volume consumed, respectively: Fig. 2B). Similarly, when RW but not HPF was present, ALMO affected neither appetitive nor consummatory processes related to RW (RW area occupancy and distance covered on RW, respectively: Fig. 2C), despite dramatically reducing the RW behaviors when both RW and HPF were available (see Fig 1G-I). These findings are incompatible with orexin-dependent modulation of isolate RW or HPF-directed behaviors as explanations for TRVE.

An alternative hypothesis is that HONs shape value comparisons between RW and HPF (Fig. 2D), which predicts that ALMO would affect RW and HPF usage only in a choice scenario (i.e. when both are available). To look at behavioral components underlying such valuation, we first examined how mice decided to enter either the RW or HPF-arm from a neutral zone (center-area). We found that control (vehicle-injected) mice are much more likely to enter the RW-arm than the HPF-arm, and this effect was abolished under ALMO (Fig 2E). These results suggest that HONs may facilitate decision-making directing the mice towards the RW over the HPF. However, given that HPF/RW entry-probabilities are equal under ALMO (Fig. 2E), what then explains the ALMO-diminished RW behaviors observed in Fig. 1? In our scenario, value-based choices not only include the decision to enter an arm, but also the decision to leave. When quantifying visit-durations to both arms, we found that ALMO significantly increased the average visit duration to the HPF when the RW was available, but had no effect once the RW was removed (Fig. 2F). ALMO did not modulate visit-durations to the RW, regardless if HPF was present or not (Fig. 2G). Together, these results suggest that HONs drive TRVE by de-homogenizing directed entry-behavior and decreasing HPF visit-duration when a RW opportunity is available.

Consummatory behaviors typically involve value-comparisons. To more directly probe the effects of HONs on the consummatory decision-making between HPF and RW, we placed mice into an environment in which both RW and HPF were available in close proximity, to minimize the influence of appetitive place-preference driven behaviors. When comparing the volume and distance travelled, we found that ALMO and vehicle-injected mice had significantly different regression coefficients (Fig 2H, Chow-test p<0.001). Control mice showed no significant relationship between running and eating, while in ALMO-injected mice eating was inversely proportional to running (Fig. 2G), thereby mice who ran less consumed increasing amounts of HPF (Pearson’s r = -0.673, p<0.001). Given the limited experimental duration, RW-commitment would presumably come at the opportunity-cost of HPF-usage, and vice versa. A strong negative relationship between HPF and RW-usage suggests an elastic demand between the two options, whereas a flatter line reveals a more inelastic, fixed demand for HPF agnostic to RW-usage. HONs therefore may drive inelastic valuations for how mice choose to spend their time between feeding and exercise opportunities, thereby preserving TRVE.

Overall, these data indicate that HONs’ function in the acute TRVE context cannot be explained by their previously-proposed roles as enhancers of physical activity ^16, 17^, or stimulators or eating ^5, 10^, or by running-induced anorexia (Fig. 2H). Instead, the results suggest that orexin transmission arbitrates between eating and running, without influencing appetitive or consummatory drives towards either.

### Orexin neuron dynamics underlying exercise prioritizing

Does HON activity track behavioral choices during TRVE, or is it simply a tonic background signal underlying TRVE? To probe this, we performed real-time photometry recordings of HON-targeted activity indicator GCaMP6 (Fig. 3A). During the 10 min sessions used to assess prioritizing, the HON signal varied considerably, both as a function of maze location (Fig. 3B) and behavioral transitions (Fig. 3C). Distinct HON activity waveforms were seen seconds before locomotor transitions from maze center to either RW or HFP arm, or before transitions from RW or HFP arm to center, consistent with an evidence-like decision-making signal (Fig. 3E). To assess more formally the extent to whether rapid HON population dynamics reports/encodes behavioral commitments, we fit a linear mixed-effects model with HON signal as the response variable and wheel-running, non-wheel locomotion, and HPF licking as input variables (Fig. 3F). We found that HON activity could be predicted from the behavioral variables (conditional R2 = 0.334, marginal R2 = 0.314; Fig. 3G). Weight estimates show HON signal was negatively related to licking and positively related to wheel and non-wheel running speed (Fig. 3I, upper). This behavioral encoding by HONs was presumably associated with behavioral control they exert when integrated over the experimental duration. To this end, comparing effect sizes of control and HON-antagonized mice indicated that HON signals increase HFP consumption and suppress running (Fig. 3I, lower). Together, these data reveal that HONs rapidly encode individual behavioral choices underlying acute prioritization of exercise over time.

**Fig. 3.**
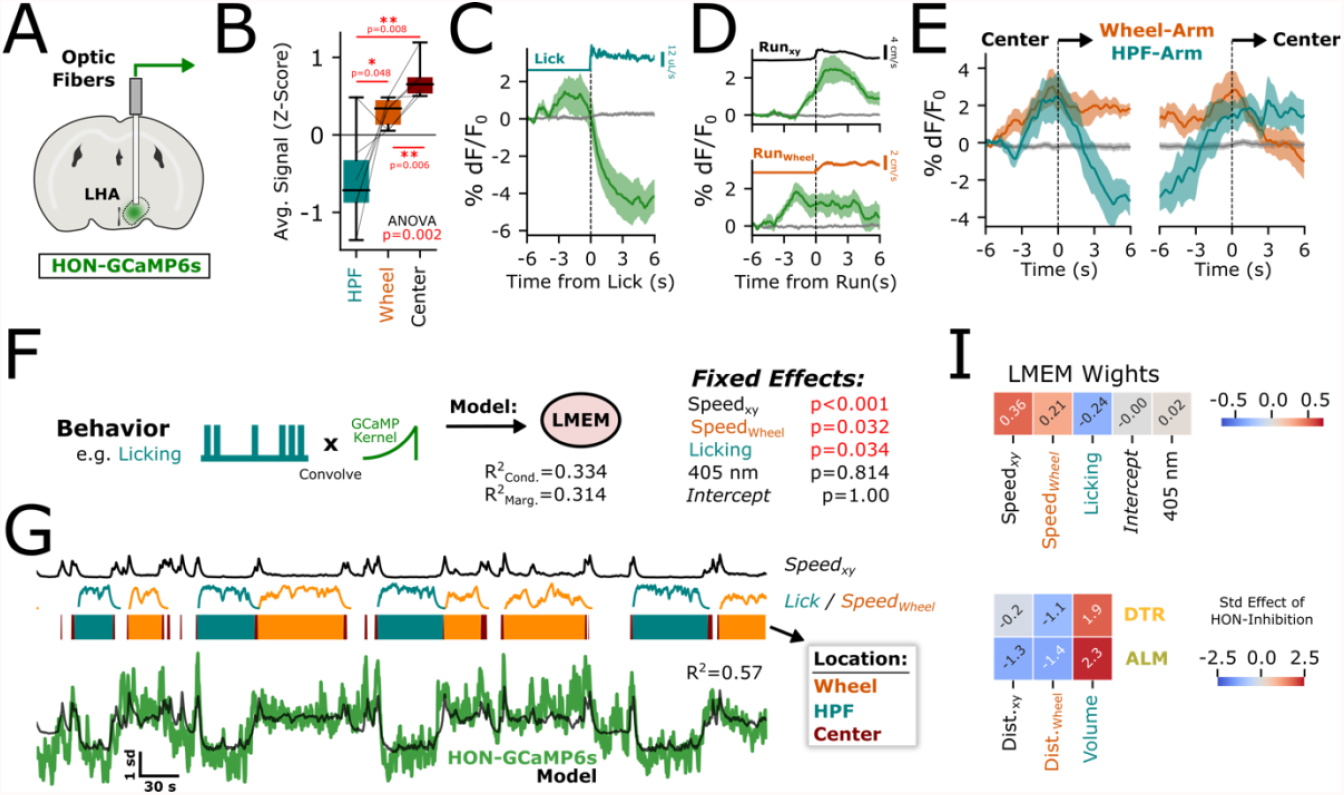
Orexin neuron activity dynamics during rapid prioritization of exercise. **A** HON activity was recorded from the LHA using GCaMP-based fiber photometry. **B** Average recorded calcium signal of n = 6 mice while in RW, HPF, or center areas. Signal was significantly different across areas, Repeated-measures ANOVA p = 0.002. Posthoc comparisons represent two-sided two sample t-tests. **C** Behavior-aligned calcium signal (green) and isosbestic 405 nm (grey) to the start of a HPF-licking bout. Mean ±SEM, baselined -6 to -5 seconds. **D** Behavior-aligned calcium signal (green) and isosbestic 405 nm (grey) to the start of a locomotor event in the open maze (upper) and on the running wheel (lower). Mean ±SEM, baselined -6 to -5 seconds. **E** Arm-entry-aligned calcium signal (orange, blue) and isosbestic 405 nm (grey)to either RW or HPF. Mean ±SEM, baselined -6 to -5 seconds. **F** Diagram of linear-mixed effects model (left) and output of fixed-effects. T-tests use Satterthwaite’s method. **G** Arena-locomotion, licking behavior, and RW-usage are plotted alongside arm-occupancy, HON photometry (green), and model-predicted HON activity (black). Model fit is given by R^2^ score of the predicted trace using LMEM estimator weights. **I** Estimator weights for the fixed effects of the complete fitted model (upper). Standard effects of inhibitory HON manipulations (lower).

## DISCUSSION

Overall, our results suggest that the HON system is an essential part of the neural hardware responsible for the desirable but poorly understood behavioral trait of seeking and performing voluntary exercise, despite the parallel temptation of highly-palatable food. In a complex environment with multivalent reward alternatives ^18, 19^, mimicking human daily life where reward choices (including exercise vs. feeding) are freely available, our data reveal that mice choose to maintain RW-directed appetitive and consummatory behaviors despite the presence of HPF. The effects of reducing endogenous HON activity were more complex than a valence-change towards RW or HPF; specifically, the effects of HON ablation were only/most-notable when mice were forced to split their time between the two options. This suggests that HONs implement this TRVE behavior through a valuation mechanism, rather than through modulating the drive to eat or exercise as previously proposed. Decision behaviors, such as the choice to enter the RW over the HPF arm, were non-homogenous in control mice; RW-arm was more likely to be entered than HPF from a neutral zone. HON deficiency was sufficient to disrupt this distinction, suggesting that normal HON-function may serve to differentiate entry-valence towards palatable options. Decisions to leave an arm were also examined, wherein HON deficiency was sufficient to increase the amount of time spent in the HPF, but not RW-arms, when both options were present. Importantly, in our experiments mice were constrained by a fixed experimental duration, whereby RW-commitment comes at the opportunity-cost of HPF-usage, and vice versa. The strong negative relationship observed between HPF and RW-usage (see Fig. 2H) suggests an elastic demand between the two options, whereas a flat line reveals an inelastic, fixed, demand for HPF agnostic to RW-usage. HONs therefore may prevent a price-elasticity based valuation between feeding and exercise opportunities, thereby preserving TRVE. At the level of neural representation of this decision-making, rapid HON activity displayed distinct action-specific signatures. A model incorporating these rapid fluctuations fit to positive-weighted estimates for locomotor-related behaviors, and negative-weighted estimates for HPF licking-behaviors. When integrated over time, these influences may explain the effects of HON loss of function manipulations. In summary, by identifying HONs are reporters and controllers of decision-making related to exercising, our data pinpoint a genetically-defined entry-point for further study of biological underpinnings of temptation-resistant voluntary exercise. The free choice paradigm established in this study is expected to aid in uncovering the underlying mechanisms that contribute to pathological imbalances between motivation for exercise or feeding. It will enable researchers to investigate whether these imbalances tend to favor feeding over running (such as in obesity) or running over feeding (such as in restrictive anorexia nervosa in extreme cases, or for a healthy lifestyle). This, in turn, will facilitate future discoveries in this field.

## METHODS

### Experimental subjects

All animal experiments followed Swiss Federal Food Safety and Veterinary Office Welfare Ordinance (TSchV 455.1, approved by the Zurich Cantonal Veterinary Office). Adult C57BL6 male mice were studied, except for in Figure 2G, where female mice were also used, thusly whether all conclusions apply to female mice remains to be determined. For HON ablation experiments (Supplemental Figure 1 and 2) we used a previously validated diphtheria (HON-DTR) ablation model ^14^. Animals were housed in reversed light-dark cycle (lights off at 7:00 AM). All experiments were performed during the dark phase. Animals had ad libitum access to water, to ensure stable motivation, mice were subjected to a mild overnight food restriction (light cycle) before behavioral experiments unless stated otherwise. When relevant, cohorts were structured to allow a pseudo-randomized crossover design.

### Surgeries and viral vectors

Activity of HONs was measured using a previously-validated orexin promotor (hORX)-driven GCaMP6s sensor. Briefly, mice were anaesthetized using 2-5% isoflurane following operative analgesia using buprenorphine and site-specific lidocaine. The GCaMP6s calcium indicator was stereotaxically injected unilaterally (randomized) into the LH (AAV1-hORX-GCaMP6s.hGH, 10^13-10^14GC/mL, Vigene Biosciences). Coordinate injections: anteroposterior [AP], -1.35; mediolateral [ML], ± 0.90; dorsoventral [DV], -5.70, -5.40, &-5.10, 70 nl at 1nl/s per site; NanoInject III injector). Optic fiber cannulas (200 μm diameter, 0.39 numerical aperture (NA) fiber with 1.25 mm ceramic ferrule; Thorlabs) were implanted unilaterally above the injection site in the LH (AP, -1.35; ML, ± 0.90; DV, - 5.00). Mice were given postoperative analgesia and allowed to recover for at least two weeks before experiments began.

### Fiber photometry and modeling

Fiber photometry experiments (Figure 3) used a multifiber camera-based photometry system (Doric). Alternating illumination at 405 nm and 465 nm at 20 Hz, average power = 70 μW. HON-GCaMP6s emission fluorescence was recorded wherein the 405 nm LED was used as an isosbestic control for movement-related artefacts, and 465 nm represented GCaMP6s calcium-dependent HON dynamics. GCaMP6s bleaching was controlled for by fitting and subtracting a triple-exponential curve to the full trace. For following analyses, each trace was either z-score normalized to the entire trace, or a locally baselined percent dF/F_0_ was used as specified.

For Fig. 3F-I, a linear mixed-effects model (LMEM) was used to predict z-scored GCaMP6s photometry signal. Licking, wheel-running speed, and speed in the xy plane were convolved (using a 60s long decay kernel with a decay rate equivalent to the reported GCaMP6s –half-life (1.796 s) ^20^ and then z-scored along with the isosbestic point (405nm) to form the fixed effects. Each mouse was a random effect with a free slope in respect to licking, wheel-running, and speed in the xy plane. 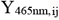 in the LMEM denotes the predicted z-scored response of the i^th^ sample from the j^th^ mouse given the fitted input variables.

The model:

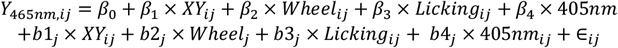

Where

- β0 is the fixed-effect intercept
- β_1-4_ are the fixed-effect slopes
- b1-4_j_ are the random-effect slopes
- ε_ij_ is the residual error

Was fitted using the lme4 library in R. We assume that all random effects are normally distributed. R2 values were computed using Nakagawa’s R2 for mixed models.

Standard effects (Cohen’s d) as in Fig. 3I were calculated as follows:

For paired samples (ALMO cohort):

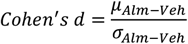

Where

- μ is the mean difference between drug and vehicle injections
- σ is the standard deviation of the difference between drug and vehicle injections

For unpaired samples (DTR cohort):

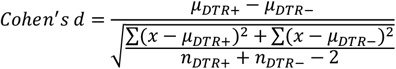

Where

- μ is the mean of the cohort
- n is the number of mice in each cohort

### A free-choice 8-arm radial maze

The subjects were introduced to a radial arm maze, featuring a central area from which eight identical arms extended outward. In our study, we modified the traditional use of the radial arm maze, originally employed by Olton and Samuelson to assess spatial learning and memory ^21^. Instead, we adapted the task to capitalize on the innate exploratory behavior of mice, allowing them to naturally engage in various activities and develop their preferences. Unlike traditional maze training paradigms, mice in this task did not require prior training to navigate and explore the maze; they exhibited spontaneous exploration from the initial testing session. Subjects were allowed to explore freely for 10 minutes. The short experimental duration was chosen since it allowed us to efficaciously study the acute, free-choice explorative strategies used by the subject. The arms contained different alternatives as follows: a running wheel (Scurry Tethered Mouse Wheel 80840WB, Lafayette Instrument Co.), normal laboratory-chow (3430 Kliba Nafag, Kaiseraugst, Switzerland), a novel same-gender conspecific mouse, a dish of water, a novel object, an illuminated “light” arm, and a dark arm. Light and dark arms were insulated with a non-transparent plastic as to minimize light spillage. A final arm was left empty at the beginning of the experiment, but was later outfitted with a custom-built HPF-dispenser (161K011, NResearch) with a capacitor-based lick-sensor (AT42QT-1010, SparkFun Electronics) to gauge consumption. When included, 6uL of HPF was dispensed for every 10 detected licks. We used milkshake (Energy milk, strawberry flavor, Emmi Schweiz AG) as HPF, since there is abundant evidence that it is highly palatable and attractive for both mice and humans ^22-26^.

To habituate the mice to the maze, on the first day mice were placed for 10 minutes into the empty maze. Then, on the following 7 days, all arm-contents (except HPF) were introduced while the position and RW activity of the mice were recorded via a ceiling-mounted camera. Only the final day was used for statistical analysis. Proper habituation is crucial to minimize stress and anxiety that could otherwise interfere with the interpretation of task performance.

Following RW-only experiments, mice were habituated to HPF in their home-cages overnight. On the following day, to ensure stable licking behavior mice were habituated to lick from a spout in a chamber with a lick-port until at least 100 rewards were obtained or 30 minutes had passed. Mice who did not learn to lick on the first day were repeated until the criterion was reached. HPF + RW experiments were performed in the radial maze. Finally, the wheel-arm was closed in entirety and mice were assessed once again for place preference in the maze without access to RW.

When relevant, vehicle/drug injections were pseudo-randomized and staggered across days. Fiber photometry was always recorded when applicable.

### Simple 2 choice maze

Behavior experiments were conducted in a custom made 194 mm x 181 mm x 398 mm acrylic chamber to assess consummatory decision-making between HPF and RW. Mice were introduced into the testing chamber where both the RW and HPF were accessible in close proximity, aiming to reduce the impact of appetitive place-preference-driven behaviors. On one side of the chamber a RW was attached and on the other side HPF was delivered in 6 μL increments via the same mechanism as in the 8-arm radial maze. While performing the task, mice moved freely in the chamber and could either choose HPF or RW for 10 min. All training sessions were conducted in the dark or under red light conditions. Mice were food deprived for 2h prior to behavioral testing to reduce variability in investigatory parameters across animals.

### Pharmacological experiments

30 mg kg^-1^ of dual orexin receptor antagonist ALMO (Almorexant (hydrochloride), MedChem Express) was dissolved in 2% dimethylsulfoxide and PBS vehicle with 25% β-CD (2 hydroxypropyl-β-cyclodextrin). ALMO or vehicle solution was administered via intraperitoneal injection 40 minutes before performing behavioral experiments. Mice had been previously habituated to intraperitoneal injections of PBS before the experimental day.

### Data analysis and statistics

Locomotion on the RW, as well as licking from the HPF, were recorded at 500 Hz using custom python scripts and a digital I/O device (Doric). Spatial location in the radial maze was recorded using an infrared camera at 5 Hz, and then extracted using a custom deep-learning network trained on over 7000 manually labeled frames. Paths were smoothed using a 3-sample boxcar filter, and then aligned to nine unique ROIs representing the eight arms of the maze + center. Points in which the mouse could not be identified made up <0.1% of the data and were linearly interpolated.

Raw data processing and statistical analysis was performed using Python. Significance was defined as the following p-values: *p<0.05, **p<0.01, ***p<0.001, and ns = p>0.05. Data are presented as boxplots of the median and inter-quartile range unless specified otherwise.

## ACKNOWLEDGEMENTS

This work in funded by ETH Zürich. DB and DPR conceived the study and designed the protocol, with contributions from ALT. ALT and XL performed most of the experiments, CS contributed to experiments in Fig. 2. ALT designed and performed all data analyses. RP advised on data analyses. DB and ALT wrote the text with inputs from DPR and RP. The authors have no competing interests to declare.

## FIGURES

**Supp. Fig. 1.**
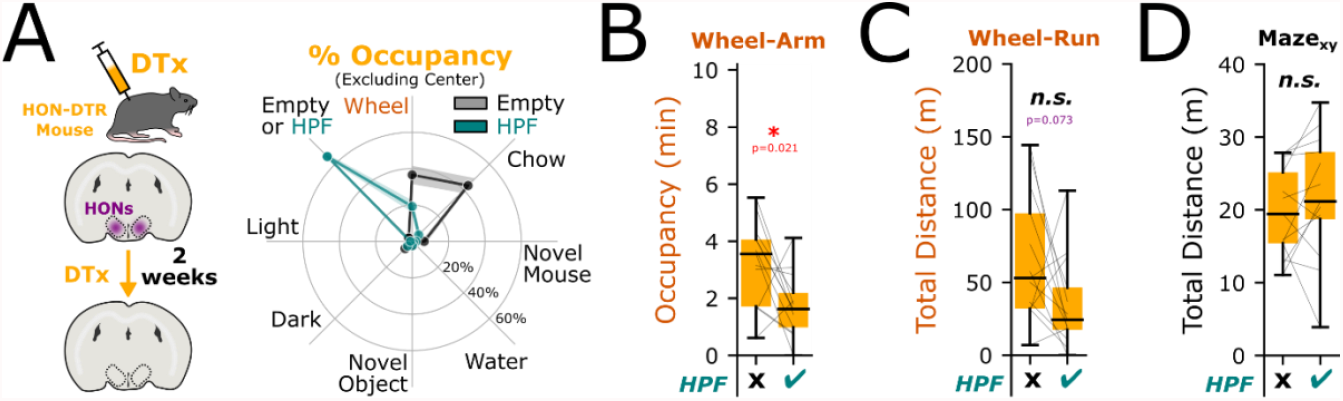
(related to Fig. 1). Effect of HON ablation on TRVE. **A** To ablate HONs, transgenic mice expressing diphtheria toxin receptor selectively in HONs (Gonzalez et al., 2016) were injected with diphtheria toxin >2 weeks before experiments began. Subjects displayed the highest % occupancy in the RW or chow-arm without HPF (grey). Subjects displayed the highest % occupancy in the RW or HPF-arm when the HPF was available (teal), note that RW-arm % occupancy was reduced when HPF was added. Lines represent mean and SEM of n=13 mice. **H** Effect of HPF on total time spent in the wheel-arm in HON-ablated subjects. Two-sided two sample paired t-test, p = 0.021. **I** Effect of HPF on total distance traveled on the RW in HON-ablated subjects. Two-sided two sample paired t-test, p = 0.073. **J** Effect of HPF on total distance run in the arena in HON-ablated subjects. Two-sided two-sample paired t-test, p = 0.280.

